# Decision-relevant measurement for precision microbiome interventions

**DOI:** 10.64898/2026.10.09.758009

**Authors:** Javad Aminian-Dehkordi, Mohammad Mofrad

## Abstract

Human gut microbiome studies often rely on stool, despite increasing evidence that it incompletely represents the spatially heterogeneous intestinal ecosystem. Whether this loss of spatial detail matters for downstream intervention decisions is largely unresolved. We treat compositional reconstruction and decision preservation as separate objectives. Using 16 matched recovery events from 11 participants, each containing stool and four intended regional capsule measurements, we aggregated amplicon sequence variants into 11 prespecified taxonomic–functional proxy guilds. We evaluated regional-to-stool models by nested leave-one-participant-out validation under complementary compositional geometries. We then assessed decision preservation through three spatial proxy states and measurement-induced decision loss (MIDL) across normalized binary intervention thresholds. Multi-region CLR ridge and Aitchison regional-plus-reference models had the lowest point-estimate Aitchison errors, but evidence that either outperformed the population Aitchison center was not retained after multiplicity correction. Rankings changed under total variation and after guild amalgamation; at the coarser resolution, the primary ordering showed a small, uncertain reversal that was sensitive to zero handling. The model with the lowest compositional error did not have the lowest MIDL-curve calibration error. Population centers closely reproduced the nearly uninformative observed-stool MIDL curves while retaining little individualized regional information. These findings show that reconstruction, decision preservation, and calibration to observed-stool decision loss answer different questions. Evaluation of microbiome measurement strategies should therefore treat compositional geometry, biological representation, and the downstream decision task as parts of the estimand.

## 1. Introduction

Designing microbiome-based interventions is fundamentally a decision problem. Researchers developing dietary strategies, probiotics, or microbiota transplantation must determine whether an intervention is warranted, which ecological feature should be targeted, and what intervention is appropriate. These decisions are difficult because gut microbial communities vary substantially among individuals, microbial associations do not necessarily identify causal or therapeutically modifiable mechanisms, and apparently similar baseline profiles can be followed by different responses to the same intervention (Kuntz and Gilbert, 2017; Gulliver et al., 2022; Zeevi et al., 2015; Zmora et al., 2018). Although microbiome-guided treatment selection is not yet routine for most diseases, precision microbiome interventions increasingly depend on measurements that distinguish biological states relevant to intervention choice.

Most human gut microbiome studies rely on stool because it is non-invasive and compatible with large-scale molecular profiling. Stool is nevertheless a downstream sample from a spatially structured ecosystem in which different factors vary along the gastrointestinal tract and create region-specific ecological niches (Donaldson et al., 2016; Tropini et al., 2017). Direct sampling under physiological conditions has shown that microbial, viral, proteomic, and metabolomic profiles can vary across regions and differ from stool (Shalon et al., 2023). Mucosal colonization and responses to administered probiotics can likewise be person-specific and only partially reflected in stool measurements (Zmora et al., 2018). Consequently, localized biological signals may be diluted, transformed during downstream transit, or weakly represented in stool. This matters when intervention decisions depend on regional processes that are not captured by broader stool-level community patterns.

The resulting problem cannot be summarized by asking whether stool and intestinal samples are compositionally similar. A large compositional difference may have little practical consequence when both measurements support the same intervention, whereas a modest difference may matter if it changes the preferred action. A model can therefore reconstruct broad features of stool composition while smoothing away a localized signal that carries decision-relevant information. This motivates two distinct empirical objectives: compositional reconstruction, which evaluates how accurately regional measurements can reproduce stool composition, and decision preservation, which evaluates whether the reconstructed signal retains the distinctions needed for a specified decision.

A structurally related problem appears in microeconomic information design, where decisions are made from imperfect observations of heterogeneous underlying states. The outcomes that can be achieved depend on which distinctions the available information preserves (Bergemann et al., 2015), and probabilistic masking can alter those attainable outcomes by changing what the decision maker can distinguish (Fallah et al., 2024). The biological setting is different, but the information structure is similar: an underlying state generates an imperfect observation, and that observation informs a downstream action (Fig. 1). Decision theory formalizes this relationship through statistical experiments and garbling: if one measurement can be generated by adding noise to another, the less informative measurement cannot be more valuable for every well-defined decision problem (Blackwell, 1953).

**Fig 1:**
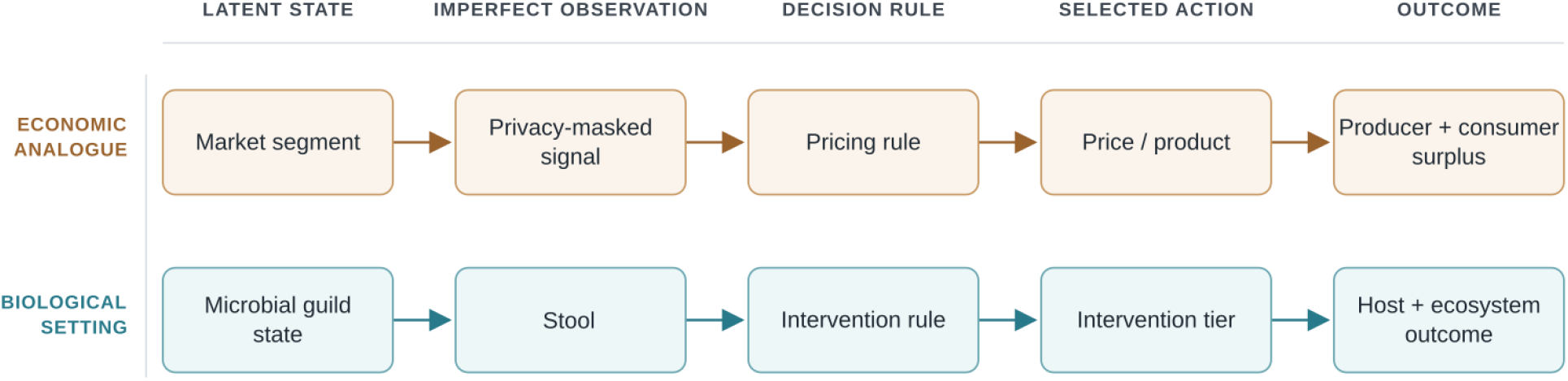
Shared information–decision architecture across economic and biological settings. Privacy masking or measurement blur converts a latent state into an imperfect observation, which a decision rule maps to an action and attainable outcome. The correspondence is structural, not semantic.

Our previous theoretical work adapted this state–measurement–decision framework to gut microbiome measurement, characterized feasible intervention outcomes under information loss, and separated measurement fidelity from decision value (Aminian-Dehkordi and Mofrad, 2026). The present study examines the empirical question left open by that framework: when regional gut measurements are used to reconstruct stool, does better compositional reconstruction also imply better preservation of information relevant to intervention decisions?

We addressed this question using matched 16S profiles from stool and four intended intestinal sampling regions reported in the literature. Taxa were aggregated into prespecified taxonomic–functional proxy guilds, and several regional-to-stool modeling strategies were evaluated under nested leave-one-participant-out validation across complementary compositional geometries. We then mapped the regional profiles to prespecified spatial proxy states and compared observed and predicted stool according to the decision loss induced across binary intervention thresholds. This design allowed us to examine if the model that best reconstructed stool composition also best preserved stool-derived decisions, how much regional measurements contributed beyond a population stool reference, and whether the conclusions were stable across alternative analytical choices. Together, these analyses test whether compositional accuracy and decision preservation identify the same measurement strategy or expose different aspects of information retained in stool.

## 2. Conceptual and mathematical framework

### 2.1. Regional compositions and stool

For participant-event *i*, let

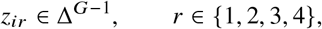

denote the guild composition measured by capsule type *r*, where Δ^*G*−1^ is the *G*-part simplex, and let *s*_*i*_ ∈ Δ^*G*−1^ denote matched stool. Capsule type indicates the intended proximal-to-distal sampling order; the exact anatomical location of an individual capsule was not directly verified in the source data. A regional-to-stool model is a map

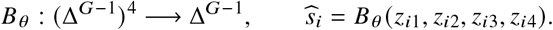

We use blurring to denote statistical compression of spatially resolved information into a single stool-like profile. The term does not imply that stool is a literal physical mixture of the sampled regions, and fitted coefficients are not interpreted as transit fractions or causal contributions.

### 2.2. Candidate blurring models

No single regional-to-stool mechanism was assumed a priori. Stool may contain little participant-specific regional information, may resemble an additive regional average, may be better represented by relative log-ratios, or may combine regional information with a strong population stool signature. We therefore compared a structured set of low-dimensional models spanning these possibilities while limiting flexibility to match the small number of participants. The complete candidate list and fitted objectives are summarized in Supplementary Table S1.

Let *ω*_*i*_ be the training-event weight, chosen so that each participant contributes equal total weight, and let 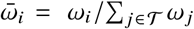. The geometry-matched no-regional-information baselines were the participant-balanced arithmetic stool center

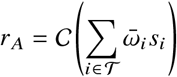

and, after multiplicative zero replacement, the Aitchison stool center

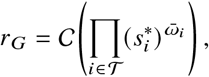

where *C* denotes closure to unit sum, 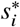 is the zero-replaced composition, and *T* is the outer-fold training set. These two centers solve different optimization problems and were retained separately: *r*_*A*_ is the natural probability-scale reference, whereas *r*_*G*_ minimizes squared Aitchison distance in log-ratio space (Aitchison, 1982; Gloor et al., 2017).

Fixed regional baselines used each capsule directly, the arithmetic mean of Capsules 3 and 4, the equal arithmetic mean of all four regions, and the equal Aitchison center of all four regions. Learned arithmetic mixtures took the form

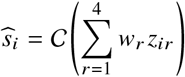

or, with population anchoring,

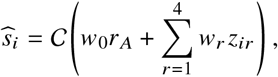

with nonnegative weights that sum to one. The weights minimized participant-balanced squared component error in the training set. Because convex combinations of closed compositions already sum to one, closure in these expressions is a numerical safeguard.

The corresponding Aitchison-geometric mixtures were defined in centered-log-ratio space:

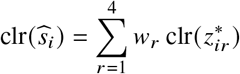

and

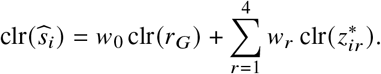

The simplex-constrained weights minimized participant-balanced squared Aitchison error. These models are weighted geometric centers; they are not conditional expectations and do not establish a physical source-mixture mechanism.

A regularized multi-region model allowed cross-guild relationships that cannot be represented by one weight per region. The four regional CLR vectors were concatenated and mapped to stool CLR coordinates as

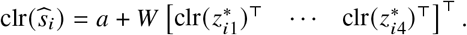

A ridge penalty 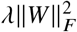 controlled the large number of coefficients relative to the cohort size, and *λ* was selected entirely within the outer training data by inner leave-one-participant-out validation (Varma and Simon, 2006).

### 2.3. Decision-preservation layer

Compositional reconstruction alone does not establish that a model preserves information relevant to an intervention. We therefore defined three prespecified binary proxy tasks that represent distinct spatial decision structures: a proximal-local oxygen-tolerant/sulfidogenic state, a distal-local low butyrate-associated state, and a Bacteroidales/Prevotella hotspot occurring in any sampled region (See SI S2.5).

For task *q*, let 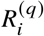 be the regional score defined in Table S3. Within each outer training fold, the intervention-relevant proxy state was

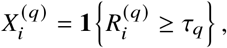

where *τ*_*q*_ was the participant-weighted training median after orienting low-state scores so that *X* = 1 always denotes the state for which intervention may be considered. For observed or model-predicted stool, a task-specific observation threshold was selected using only training participants to minimize participant-weighted state misclassification. Held-out observations produced cross-fitted signals 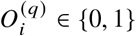 ∈ {0, 1} and an empirical binary channel

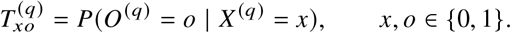

To avoid assigning an unsupported drug, dose, or disease-specific utility, we evaluated the complete normalized binary intervention-threshold class. Let *d* ∈ {0, 1} denote the intervention decision, with *d* = 1 representing the higher intervention tier. For threshold *η* ∈ [0, 1],

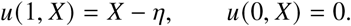

Thus, intervention yields normalized benefit 1 − *η* in state *X* = 1 and burden *η* in state *X* = 0. If *P*_*o*_ = *P*(*X* = 1 | *O* = *o*), the optimal decision rule selects the higher tier when *P*_*o*_ ≥ *η*. The expected value supported by channel *T* is

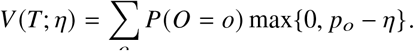

Identity observation reveals *X*, giving *V* (*I*; *η*) = ϕ_1_(1 − *η*) for ϕ_1_ = *P*(*X* = 1). Conditional decision loss is therefore

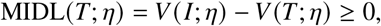

where nonnegativity follows because a decision-maker with identity information can imitate any rule based on a coarser signal (Blackwell, 1953; Aminian-Dehkordi and Mofrad, 2026). Following the terminology of our theoretical framework, we refer to this quantity as measurement-induced decision loss (MIDL). We summarized a channel by the uniform-threshold integral 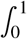 MIDL (*T*; *η*) *d*η. This dimensionless quantity averages across normalized intervention environments and is not a percentage of clinical benefit. For blurring model *B*, MIDL-curve calibration error was

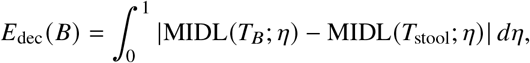

which measures how closely model-predicted stool reproduces the decision loss observed from actual stool. This calibration quantity is distinct from decision preservation itself: a constant model can match a nearly uninformative stool channel while preserving little participant-specific regional information.

For a target-state prevalence *ϕ*_1_, a completely uninformative observation has integrated decision loss

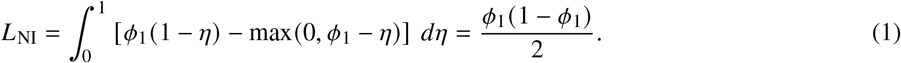

We therefore report the fraction of identity-observation decision value retained by a channel *T* as

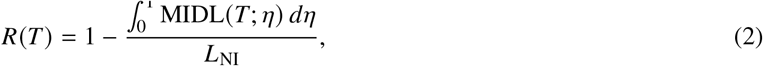

where *R* = 0 denotes a no-information signal and *R* = 1 denotes identity observation. This normalization is task-specific because the ceiling depends on ϕ_1_.

## 3. Methods

### 3.1. Data source and study scope

We performed a secondary analysis of publicly released, de-identified 16S rRNA gene sequencing exports from a study that collected matched stool and ingestible-capsule samples intended to represent four locations along the intestinal tract under physiological conditions (Shalon et al., 2023). The available files comprised processed sample metadata, an ASV count matrix, taxonomic assignments, and a data dictionary. The source study designated 16S samples for exclusion when sequencing depth was below 2,500 reads or when a sample represented the lower-priority member of a duplicate sequencing run. We applied those source-study criteria without adding an outcome-dependent filter. After filtering, all sample identifiers had to be unique and present in the metadata and ASV matrix. ASV columns were retained only when a taxonomy record was available. Taxonomic annotations at family, genus, and species levels were mapped by fixed hierarchical rules to 11 taxonomic–functional proxy guilds. A complete quality control flow and file-level hashes are provided in Supplementary Information.

### 3.2. Model fitting and nested validation

All reconstruction models described in the Conceptual and mathematical framework were evaluated by outer leave-one-participant-out validation. For each held participant, every stool center, zero-replaced reference, regional mixture weight, observation threshold, and regularization parameter was estimated from the remaining participants only. Within a training set, an event from participant *s* received weight proportional to 1/*n*_*s*_, where *n*_*s*_ was that participant’s number of training events; weights were normalized to have mean one. Thus, each participant contributed equal total weight to reference estimation and model fitting despite unequal numbers of recovery events.

Arithmetic mixture weights were estimated by constrained optimization of participant-weighted squared component error. Aitchison mixture weights minimized participant-weighted squared Euclidean distance between CLR coordinates. The probability-scale baseline was the training-fold arithmetic stool center, whereas Aitchison-space comparisons used the training-fold Aitchison stool center. Using geometry-matched centers prevents an Aitchison model from being credited for improvement that arises only because it was compared with an arithmetic reference.

### 3.3. Reconstruction endpoints

Aitchison distance was the primary reconstruction endpoint because sequencing profiles are compositions and their information is represented by relative log-ratios (Aitchison, 1982; Gloor et al., 2017)

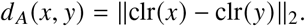

Total-variation distance, 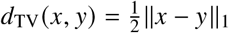,, was the main probability-scale secondary endpoint and quantifies the absolute compositional mass that would need to be reassigned. Jensen–Shannon distance was also reported. Event-level errors were averaged within participant before descriptive summaries and inferential comparisons.

### 3.4. Cross-fitted decision-layer estimation

The three task definitions in Table S3 were fixed before evaluating model performance. Within each outer fold, the regional target threshold was the participant-weighted median among training events. For actual stool and each candidate model, candidate observation cutoffs were the midpoints between consecutive unique training scores, together with the two constant-signal cutoffs. The selected cutoff minimized participant-weighted binary misclassification of the regional state. This calibration was repeated separately for each task and model because the scale of a guild score can change after compositional transformation.

After all outer folds were complete, cross-fitted held-participant states and signals were combined to estimate one population-level channel per task and model. Each participant contributed total weight one across their events. A Jeffreys-type pseudocount of 1/2 was added to each binary channel cell and to each state count to stabilize probabilities in the small cohort (Jeffreys, 1946).

### 3.5. Regional contribution and sensitivities

The regional models contain correlated inputs, so fitted simplex weights alone cannot establish that a capsule region is uniquely informative. We therefore treated the weights as descriptive and complemented them with leave-one-region-out ablation. In each outer fold, one capsule type was omitted, the reference-anchored arithmetic and Aitchison models were refit using the remaining regions, and the change in held-participant error was compared with the corresponding full model. Because the remaining regions can compensate after refitting, this analysis estimates incremental predictive contribution conditional on the model and cohort; it is not an anatomical causal effect.

Sensitivity analyses addressed three assumptions that could materially affect the conclusions. First, all events for each participant were averaged to one participant-level profile before refitting, testing whether repeated recovery events influenced the model ranking. Second, the 11 proxy guilds were collapsed to six prespecified broader ecological groups, testing dependence on taxonomic resolution. Third, because guild amalgamation changes both the retained log-ratio balances and the zero pattern, guild resolution was crossed with zero-replacement values 10^−8^, 10^−6^, and 10^−4^. For each combination, the Aitchison regional-plus-reference model was compared with its geometry-matched population center using participant-level paired differences. We also reran the decision layer at six-guild resolution; each proxy-task score was algebraically unchanged because its *K* = 6 group was the exact sum of its *K* = 11 members, while reconstruction models were refitted at the coarser representation. We verified that the selected ridge penalties did not lie at the upper boundary of the search grid. A four-state latent model, disease-specific optimizer, physical treatment dose, and clinical capsule-indication rule were not estimated because the cohort size and available outcomes do not identify those quantities. More details on the methods are provided in Supplementary Information section.

## 4. Results

### 4.1. Compositional reconstruction depended on geometry

The raw metadata contained 392 records from 15 participants. Application of the source-study 16S criteria retained 297 samples from 15 participants, all with at least 2,500 reads. The aligned ASV and taxonomy tables contained 2,437 features. 16 recovery events from 11 participants contained matched stool and all four capsule types (Figure 2); 4 participants contributed more than one complete event.

**Fig 2:**
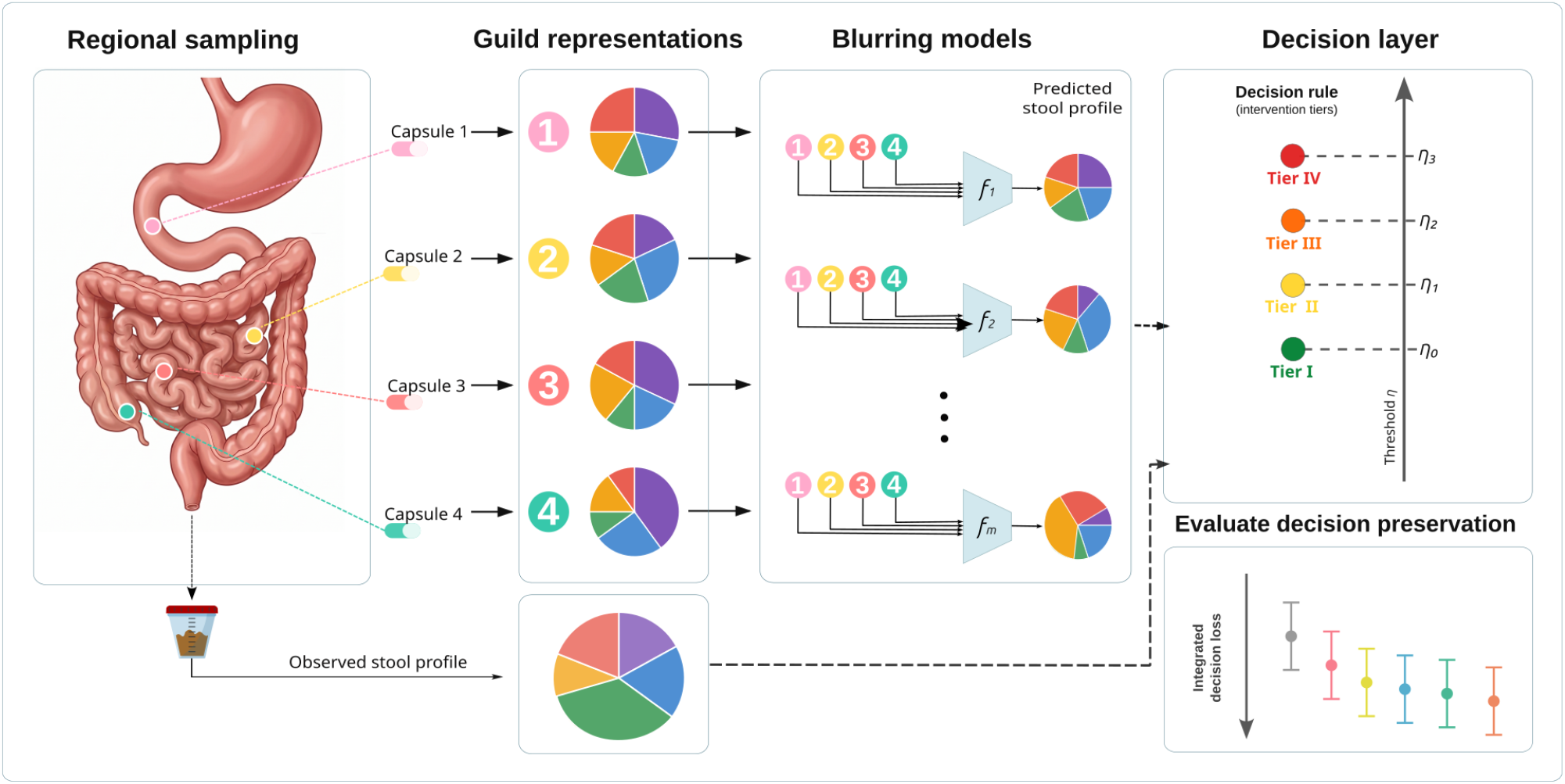
Study design and analysis architecture. Capsule types 1–4 are intended regional measurements. Compositional blurring models map all four profiles to predicted stool; a separate cross-fitted decision layer assesses preservation of three prespecified spatial proxy states.

Under the primary Aitchison endpoint, multi-region CLR ridge had the lowest point-estimate mean participant-level error (7.013 [5.593, 8.406]), followed by the Aitchison regional-plus-reference model (7.074 [6.297, 7.921]). The training-fold population Aitchison stool center had mean error 8.039 [6.582, 9.416]. Improvements over that center were 1.026 units for CLR ridge and 0.965 for the Aitchison mixture, but neither comparison remained significant after Holm correction (adjusted *P* = 0.243 and *P* = 0.243, respectively; Table 1 and Figure 3).

**Table 1:** Nested leave-one-participant-out reconstruction performance after participant-balanced fitting. Values are means of participant-level errors with 95% participant-bootstrap intervals. Lower is better.

| Model | Aitchison distance | Total-variation distance |
| --- | --- | --- |
| Population arithmetic stool center | 10.061 [8.644, 11.512] | 0.316 [0.256, 0.385] |
| Population Aitchison stool center | 8.039 [6.582, 9.416] | 0.332 [0.257, 0.417] |
| Equal arithmetic regional mean | 7.578 [6.869, 8.296] | 0.538 [0.474, 0.626] |
| Arithmetic regions + stool reference | 9.721 [8.368, 11.092] | 0.319 [0.264, 0.388] |
| Equal Aitchison regional center | 7.752 [6.950, 8.693] | 0.577 [0.510, 0.659] |
| Aitchison regions + stool reference | 7.074 [6.297, 7.921] | 0.416 [0.322, 0.516] |
| Multi-region CLR ridge | 7.013 [5.593, 8.406] | 0.343 [0.255, 0.440] |

**Fig 3:**
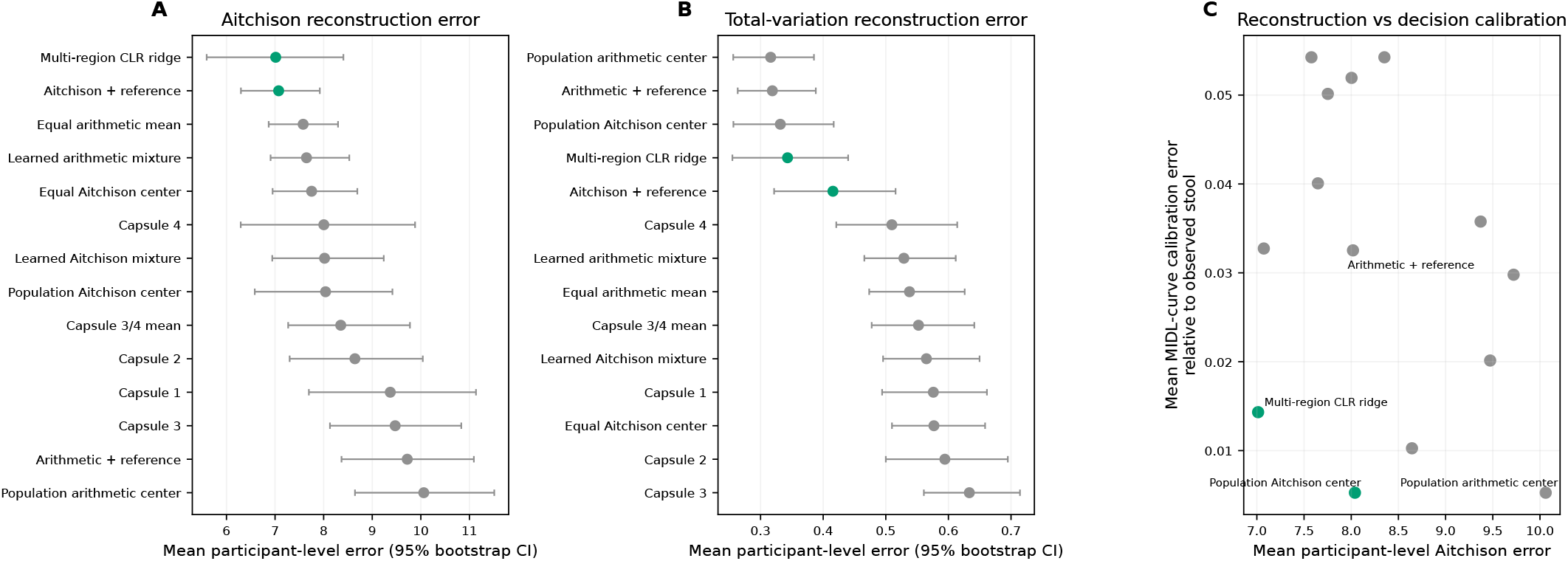
Compositional reconstruction and decision calibration. (A,B) Mean participant-level Aitchison and total-variation errors with bootstrap intervals. (C) Aitchison error versus MIDL-curve calibration error relative to observed stool. Green points are on the empirical reconstruction–decision Pareto frontier.

The ranking changed under total variation. The arithmetic regional-plus-reference model (0.319 [0.264, 0.388]) was similar to the training-fold arithmetic stool center (0.316 [0.256, 0.385]; adjusted *P* = 0.752). CLR ridge also did not differ from that center after correction, whereas the Aitchison regional-plus-reference model had higher TV error (0.416 [0.322, 0.516]; adjusted *P* = 0.026). Thus, no model was uniformly preferred across log-ratio and probability-scale geometries.

### 4.2. Population anchoring and regional contributions

The arithmetic regional-plus-reference model assigned a median weight of 0.913 to the population stool reference. In Aitchison geometry, the median reference weight was 0.538, with regional weight concentrated on Capsule 4 (0.303) and Capsule 2 (0.146) (Figure 4). These are predictive coefficients, not physical source proportions. We also perfomed leave-one-region-out refitting. Omitting Capsule 4 increased mean participant-level Aitchison error by 0.191 [−0.491, 0.776]; omitting Capsule 2 changed it by 0.102 [−0.004, 0.191]. The corresponding changes after omitting Capsules 1 and 3 were −0.060 [−0.147, −0.008] and −0.020 [−0.037, −0.005]. No ablation contrast remained significant after Holm correction.

**Fig 4:**
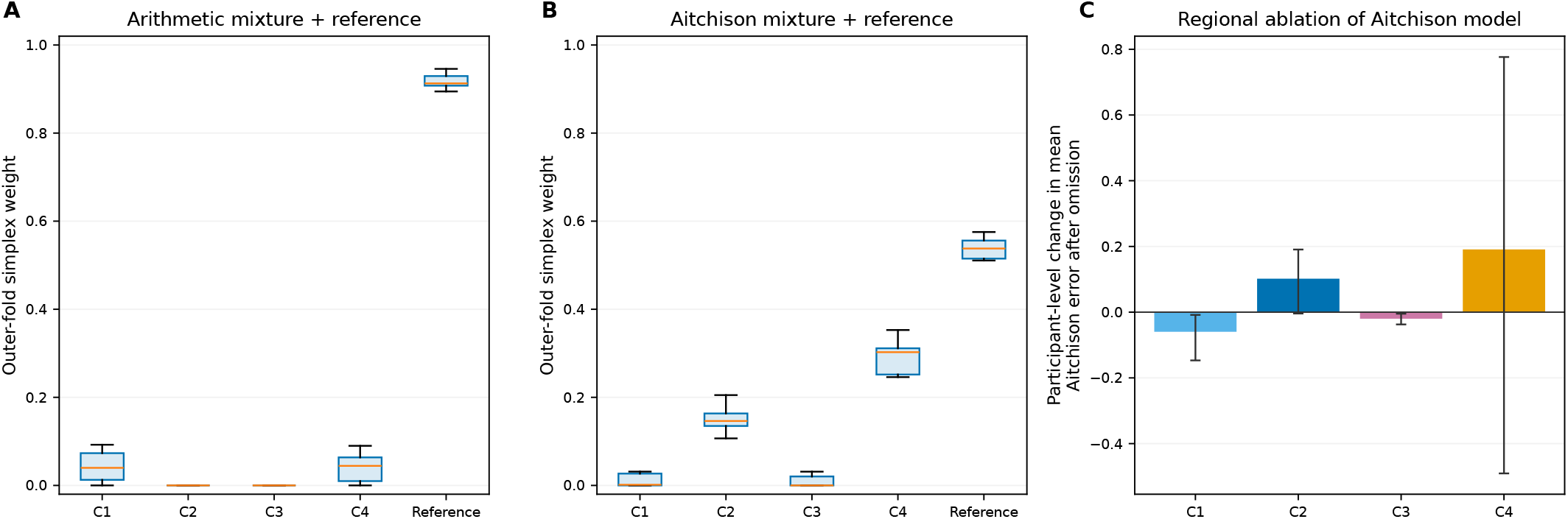
Reference anchoring and regional contribution. (A,B) Outer-fold simplex weights for arithmetic and Aitchison reference-anchored models. (C) Participant-level change in mean Aitchison error after omitting each capsule and refitting the Aitchison model; bars show means and 95% participant-bootstrap intervals. Positive values indicate worse prediction after omission.

### 4.3. Compositional reconstruction and decision preservation

Observed stool retained nonzero MIDL for all three proxy tasks. Participant-bootstrap MIDL was 0.104 [0.056, 0.124] for the proximal-local proxy, 0.113 [0.073, 0.124] for the distal-local low butyrate-associated proxy, and 0.111 [0.064, 0.124] for the regional hotspot proxy (Figures 5 and 6). These dimensionless summaries average MIDL uniformly over *η* ∈[0, 1] and are not clinical effect sizes.

**Fig 5:**
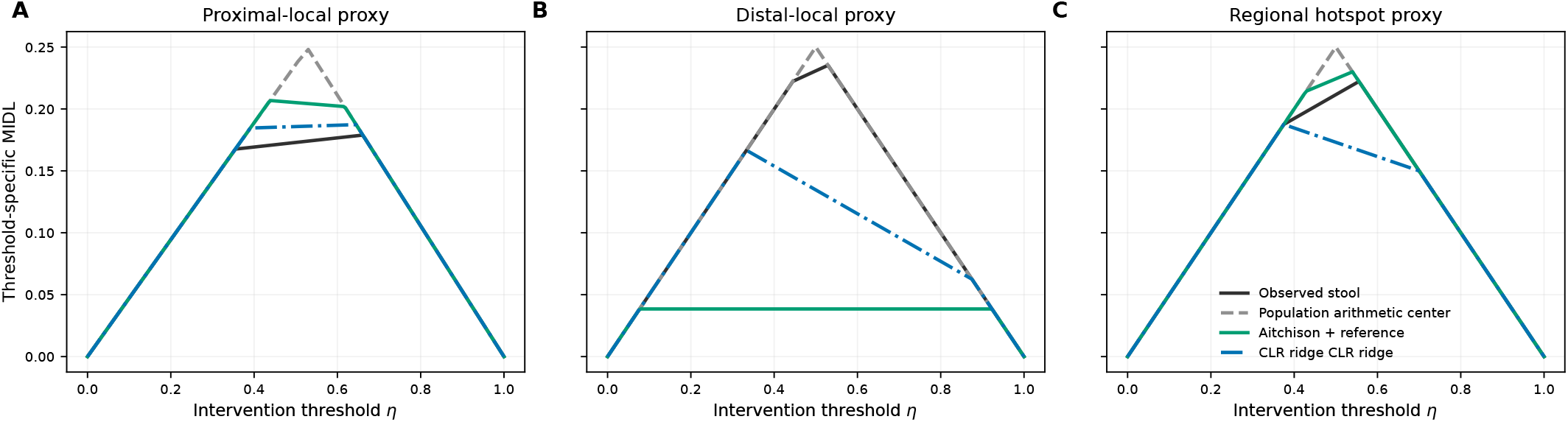
Threshold-specific MIDL curves for observed stool and selected reconstruction models. Curves compare observed stool with the Aitchison regional-plus-reference model and multi-region CLR ridge for each prespecified spatial proxy task.

**Fig 6:**
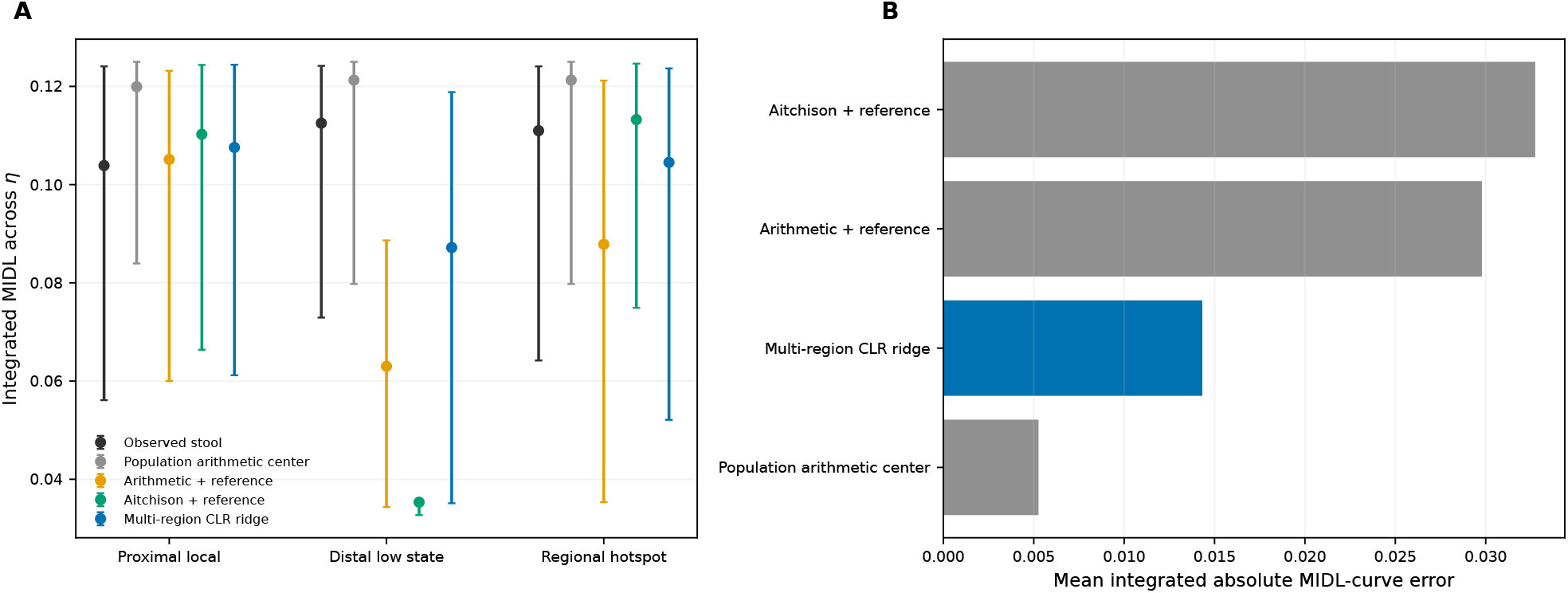
Decision preservation and calibration. (A) Participant-bootstrap intervals for integrated MIDL. (B) Mean integrated absolute deviation between model-derived and observed-stool MIDL curves.

**Table 2:** Decision-preservation and calibration results. Integrated MIDL is summarized by the participant-bootstrap median and 95% interval. MIDL-curve calibration error is the integrated absolute difference from the observed-stool MIDL curve.

| Proxy task | Measurement/model | Integrated loss | Decision value retained | Curve error |
| --- | --- | --- | --- | --- |
| Proximal-local oxygen-tolerant/ sulfidogenic proxy | Observed stool | 0.105 [0.055, 0.124] | 10.0% [0.0, 48.4] | — |
|  | Population arithmetic stool center | 0.121 [0.084, 0.125] | 0.2% [0.0, 2.9] | 0.0115 |
|  | Population Aitchison stool center | 0.121 [0.084, 0.125] | 0.2% [0.0, 2.9] | 0.0115 |
|  | Arithmetic regions + stool reference | 0.107 [0.062, 0.124] | 10.1% [0.0, 38.9] | 0.0070 |
|  | Aitchison regions + stool reference | 0.111 [0.063, 0.124] | 4.8% [0.0, 41.3] | 0.0075 |
|  | Multi-region CLR ridge | 0.109 [0.062, 0.124] | 6.6% [0.0, 41.4] | 0.0035 |
| Distal-local low butyrate-associated proxy | Observed stool | 0.113 [0.074, 0.124] | 3.1% [0.0, 30.6] | — |
|  | Population arithmetic stool center | 0.121 [0.080, 0.125] | 0.2% [0.0, 3.3] | 0.0008 |
|  | Population Aitchison stool center | 0.121 [0.080, 0.125] | 0.2% [0.0, 3.3] | 0.0008 |
|  | Arithmetic regions + stool reference | 0.063 [0.034, 0.094] | 47.9% [19.5, 71.4] | 0.0591 |
|  | Aitchison regions + stool reference | 0.035 [0.033, 0.035] | 70.9% [60.3, 71.6] | 0.0887 |
|  | Multi-region CLR ridge | 0.087 [0.035, 0.119] | 25.0% [1.4, 66.8] | 0.0304 |
| Regional Bacteroidales/ Prevotella hotspot proxy | Observed stool | 0.111 [0.064, 0.124] | 4.5% [0.0, 34.3] | — |
|  | Population arithmetic stool center | 0.121 [0.080, 0.125] | 0.2% [0.0, 3.3] | 0.0035 |
|  | Population Aitchison stool center | 0.121 [0.080, 0.125] | 0.2% [0.0, 3.3] | 0.0035 |
|  | Arithmetic regions + stool reference | 0.088 [0.046, 0.121] | 21.4% [0.4, 59.5] | 0.0233 |
|  | Aitchison regions + stool reference | 0.114 [0.076, 0.124] | 2.6% [0.0, 23.8] | 0.0020 |
|  | Multi-region CLR ridge | 0.103 [0.054, 0.124] | 10.0% [0.0, 49.3] | 0.0090 |

At point estimate, the two population stool centers had the smallest mean MIDL-curve calibration error relative to observed stool (0.0053 across tasks), followed by multi-region CLR ridge (0.0143). The arithmetic and Aitchison regional-plus-reference models had mean calibration errors of 0.0298 and 0.0327, respectively. The largest discrepancy occurred for the distal-local low butyrate-associated task. The Aitchison model had median integrated loss 0.035 and differed from observed stool by −0.077 [−0.089, −0.039]; the arithmetic reference-anchored model differed by −0.046 [−0.082, −0.011]. Both negative contrasts indicate that these models preserved more distal-local signal than actual stool. The corresponding CLR-ridge contrast was −0.022 [−0.069, 0.017], with an interval that included zero. For the proximal-local and regional-hotspot tasks, bootstrap contrast intervals for all three fitted multiregion models included zero. Improved compositional reconstruction therefore did not, by itself, establish faithful reproduction of stool-guided decisions.

Because the two local proxy states were defined from one capsule, we repeated their decision analyses after excluding that target-defining region from model fitting and prediction. For the distal-local task, the Aitchison regional-plus-reference model’s point-estimate integrated MIDL increased from 0.0355 to 0.0651 after Capsule 4 was omitted. This attenuation indicates that part of the apparent preservation arose from direct access to the state-defining region. The remaining signal is descriptive and may reflect correlation among regions; it is not an independent validation (Supplementary Table S4).

### 4.4. Sensitivity analyses

Sensitivity analyses evaluated whether the principal reconstruction conclusions depended on repeated events, guild resolution, zero replacement, or ridge regularization. Averaging repeated events to one profile per participant preserved the primary pattern: the regional-plus-reference model had lower point-estimate Aitchison error than the population center (7.025 versus 8.051). At *K* = 11 and *δ* = 10^−6^, the paired mean difference between the population center and regional model was 0.965 [−0.063, 1.892] Aitchison units (positive favors the regional model; exact *P* = 0.095). At *K* = 6, the corresponding contrast was −0.077 [−0.322, 0.153] (exact *P* = 0.560), with mean errors 3.836 for the population center and 3.914 for the regional model. The *K* = 6 reversal was therefore small and uncertain, not evidence that the population center was definitively superior.

The crossed sensitivity showed that guild resolution and zero replacement interacted. At *K* = 11, the regional model remained below its matched center across all three replacement values, with relative error reductions from 7.93% to 12.82%. At *K* = 6, the relative reductions were −7.88%, −2.01%, and 1.96% at 10^−8^, 10^−6^, and 10^−4^, respectively. Thus, the direction of the small K=6 contrast changed with zero handling.

At *K* = 6, proxy-state and observed-stool scores were algebraically identical to their K=11 definitions, while reconstruction models were refitted on the amalgamated profiles. CLR ridge then had the smallest point-estimate mean MIDL-curve calibration error (0.003), followed by the population center (0.0053) and Aitchison regional model (0.0144). These descriptive results show that representation affected both reconstruction and decision calibration; they do not replace the primary *K* = 11 analysis. The ridge optimum remained interior at *λ* = 100.

## 5. Discussion

This study evaluated regional-to-stool modeling as a two-objective problem: reconstructing a matched stool composition and preserving the regional distinctions used by a downstream intervention decision rule. The analyses yielded principal findings. Selected regional models had lower point-estimate Aitchison error, but improvements were uncertain; probability-scale rankings differed; and the model with the lowest compositional error did not have the lowest MIDL-curve calibration error.

MIDL and MIDL-curve calibration also require separate interpretation. MIDL quantifies information forgone relative to identity observation for a specified task. By contrast, *E*_dec_ examines if predicted stool reproduces the MIDL curve of observed stool. A constant population center can have low *E*_dec_ when observed stool is nearly uninformative, even though the center preserves almost no individualized regional information. Low reconstruction error, low MIDL, and low MIDL-curve calibration error therefore answer different scientific questions.

Guild coarsening was not merely a technical tuning exercise. Amalgamating 11 proxy guilds into six broader groups removed within-group balances and changed which biological distinctions contributed to Aitchison distance and model fitting. The within-representation point-estimate ordering changed, showing that the apparent value of regional information was conditional on the representation. However, the K=6 contrast was uncertain and changed direction across zero-replacement values. The supported conclusion is representation dependence and ranking instability, not that a coarser representation makes the population center superior. Measurement quality is conditional on the biological features retained, the comparison geometry, and the downstream task.

This distinction connects the empirical findings to the information-design framework developed in our previous work (Aminian-Dehkordi and Mofrad, 2026). A composition can be close to stool globally while crossing a task-specific threshold differently, and a compositionally imperfect model can preserve the same intervention tier. The three tasks here are transparent stress tests based on spatial taxonomic proxies, not validated disease biomarkers or treatment-response rules.

The framework has a direct implication for future intervention studies. A prospective dataset containing paired regional and stool measurements, a prespecified disease or response state, and observed treatment outcomes could replace the proxy states with validated targets and could attach evidence-based costs to intervention errors. Such data would allow estimation of treatment-specific decision value and of when an additional regional measurement is worth its burden. The present healthy observational cohort cannot identify a medicine, a physical dose, or a patient-specific capsule-indication rule, and none is inferred here.

### Limitations

The complete-event analysis included 16 events from 11 healthy participants, and some recovery events averaged multiple swallowed sets. Participant-balanced fitting, held-participant validation, participant-level tests, and clustered bootstrap inference reduce pseudoreplication but do not create external validation. Capsule types reflect intended regional ordering; exact anatomical positions were not directly confirmed. The profiles were 16S-derived taxonomic proxies, not direct measurements of metabolic flux, disease status, mucosal communities, or treatment response. Guild definitions and the three state maps are biologically motivated but not clinically validated. Log-ratio methods require zero replacement, and absolute Aitchison values changed with that numerical choice. The candidate mixtures are statistical mappings; their weights are not mechanistic transit coefficients. Decision thresholds were optimized within a small training cohort and then cross-fitted, but channel estimates remained sparse and were stabilized by a Jeffreys-type pseudocount. Finally, all model selection and sensitivity analyses used one dataset, so the findings require prospective validation in cohorts with paired regional, stool, and intervention-response measurements.

## 6. Conclusion

Four regional intestinal compositions contained information that could be used to construct stool-like profiles, but the analysis did not identify a universal regional-to-stool blurring mechanism. Findings showed that rankings depended on compositional geometry and guild representation, while MIDL analysis showed that reconstruction quality did not determine decision preservation or calibration to observed-stool decision loss. Microbiome measurements intended for intervention should therefore be evaluated separately for compositional fidelity, retained decision value, and decision calibration, with biological representation and task treated as parts of the estimand.

## Data and code availability

This study analyzed de-identified 16S rRNA gene-sequencing data originally reported by Shalon et al. Shalon et al. (2023). The underlying sequencing reads are publicly available from the NCBI Sequence Read Archive under BioProject accession PRJNA822660. The processed abundance tables, taxonomy data, and de-identified sample metadata used in the present analyses are available at https://github.com/mofradlab/midloss.

## Competing Interests

The authors have declared that no competing interests exist.

## Supplementary Information

### S1. Theoretical assumptions used by the empirical decision layer

The empirical decision layer requires only finite states, observations, intervention tiers, and a specified utility class. It does not require stool to be a scalar affine blur. The nonnegativity of MIDL follows because identity observation reveals *X* and can imitate any decision rule based on a coarser observation. For a finite binary channel, *V* (*T*; *η*) is the maximum of affine functions of *η* and is piecewise linear; therefore MIDL(*T*; *η*) is also piecewise linear. No claim of globally linear loss in a scalar blurring parameter is used. The feasible-set and Blackwell results are developed more generally in our previous theoretical work (Aminian-Dehkordi and Mofrad, 2026).

### S2. Supplementary methods and results

#### S2.1. Data-processing and quality-control details

The processed exports contained 392 metadata records from 15 participants. Restriction to the primary study series and application of the source-study sequencing and duplicate-run exclusions retained 297 16S profiles: 58 stool, 56 Capsule 1, 54 Capsule 2, 55 Capsule 3, 45 Capsule 4, and 29 saliva samples. All retained profiles had at least 2,500 reads. Sample identifiers were intersected across metadata, count, and taxonomy files before aggregation; 2,437 ASVs had aligned taxonomic records. The complete-event requirement yielded 16 matched stool–four-capsule events from 11 participants. When several swallowed sets were associated with the same stool recovery time, replicate profiles of the same capsule type were averaged at the guild-composition level. 4 participants contributed repeated complete events. The full sample-flow table, ASV-to-proxy mapping, event table, and input hashes are distributed with the code.

#### S2.2. Guild definitions and six-group amalgamation

The K=6 representation was defined by summing K=11 guilds before closure and model fitting. This changes the biological balances retained by the composition; it is not a change in formal taxonomic rank. K6 groups G1–G6 denote proximal/sulfidogenic, Bacteroidales/Prevotella, lactate/propionate, fiber/butyrate, mucin/barrier, and other/uncertain amalgamations, respectively.

| K=11 guild | Interpretation | K=6 amalgamation |
| --- | --- | --- |
| G01 | Sulfidogenic, bile-associated, proteolytic, or pathobiont-enriched taxa. | K6 G1 |
| G02 | Facultative, oxygen-tolerant, oral/proximal taxa often underrepresented in distal stool. | K6 G1 |
| G03 | Bacteroidales generalists with broad polysaccharide/protein versatility. | K6 G2 |
| G04 | Prevotella/Prevotellaceae carbohydrate-associated propionate proxy. | K6 G2 |
| G05 | Lactate/succinate-utilizing and propionate-associated anaerobes. | K6 G3 |
| G06 | Bifidobacteria acetate/lactate producers and cross-feeding partners. | K6 G3 |
| G07 | Ruminococcaceae/Faecalibacterium-like fiber and butyrate-associated taxa. | K6 G4 |
| G08 | Mucin-glycan and mucus/barrier-associated taxa. | K6 G5 |
| G09 | Lachnospiraceae-type butyrate/cross-feeding anaerobes. | K6 G4 |
| G10 | Other anaerobic fermenters not specific enough for higher-confidence assignment. | K6 G6 |
| G11 | Unclassified or low-information taxa retained for compositional closure. | K6 G6 |

#### S2.3. Observation-threshold implementation

For a training score vector *y* with unique ordered values *y* _(1)_ < · · · < *y* _(*m*)_, candidate cutoffs were −∞, +∞, and (*y* (_*j*_) + *y* (_*j*_+ _1_))/2 for *j* = 1, …, *m*−1. This enumerates every distinct binary partition obtainable by a one-dimensional threshold. Cutoffs were ranked lexicographically by participant-weighted misclassification error, absolute difference between predicted and target high-state prevalence, distance from the participant-weighted median score, and the numeric cutoff. This deterministic rule was used only within the training portion of each outer fold.

#### S2.4. Complete candidate-model list

**Table S2:** All candidate models and their role.

| Model | Definition and interpretation |
| --- | --- |
| Population arithmetic stool center | Participant-balanced training-fold arithmetic center; no held-participant regional information. |
| Population Aitchison stool center | Participant-balanced training-fold geometric center after zero replacement; geometry-matched no-regional-information baseline for Aitchison error. |
| Capsules 1–4 | Each regional profile used directly as a stool proxy. |
| Capsule 3/4 mean | Arithmetic mean of the two intended distal capsule types. |
| Equal arithmetic mean | Equal four-region convex average in the simplex. |
| Learned arithmetic mixture | Participant-balanced nonnegative regional weights summing to one, optimized by component-scale squared error. |
| Arithmetic + reference | Learned convex combination of four regions and the arithmetic stool center. |
| Equal Aitchison center | Equal geometric center of the four regional compositions in CLR space. |
| Learned Aitchison mixture | Participant-balanced simplex-constrained regional weights optimized by squared Aitchison error. |
| Aitchison + reference | Weighted geometric center of four regions and the Aitchison stool center. |
| Multi-region CLR ridge | Participant-balanced regularized multivariate CLR regression using all four regional CLR vectors; penalty selected inside each outer fold. |

#### S2.5. Prespecified spatial proxy tasks

The definitions use only taxonomic groups measured in the available 16S data and do not assert a diagnosed disease, measured metabolic function, or validated treatment response.

**Table S3:** Prespecified spatial proxy tasks used in the decision-preservation analysis.

| Proxy task | Regional state definition |  |
| --- | --- | --- |
| Proximal-local oxygen-tolerant/sulfidogenic state | High summed relative abundance in Capsule 1 | Sulfidogenic, bile-associated, proteolytic, or pathobiont-enriched taxa, including |
| Distal-local low butyrate-associated state | Low summed relative abundance in Capsule 4 | Ruminococcaceae-, *Faecalibacterium*- |
| Regional Bacteroidales/Prevotella hotspot | High maximum summed score across Capsules 1–4 | Bacteroid |

The remaining curated categories—lactate-utilizing or propionate-associated taxa, *Bifidobacterium*-associated acetate/lactate producers, mucin-associated taxa, and unresolved or low-information taxa—were not used to generate additional tasks. This restriction was set to limit multiplicity and to evaluate three distinct spatial summaries. Their exclusion does not imply biological irrelevance; the complete mapping rubric is provided in the Supplement and reproducibility files.

#### S2.6. Target-region-omission decision analysis

The target state remained defined from the prespecified local region, but that region was excluded from model fitting and prediction. The hotspot task has no single fixed target region and was not included.

**Table S4:** Point-estimate decision results after omitting the target-defining region from regional-to-stool model fitting and prediction.

| Task | Omitted | Model | Integrated loss | Retained | Agreement |
| --- | --- | --- | --- | --- | --- |
| Proximal-local oxygen-tolerant/sulfidogenic proxy | Capsule 1 | Arithmetic + reference, target region omitted | 0.1053 | 15.5% | 0.688 |
| Proximal-local oxygen-tolerant/sulfidogenic proxy | Capsule 1 | Aitchison + reference, target region omitted | 0.1206 | 3.2% | 0.688 |
| Proximal-local oxygen-tolerant/sulfidogenic proxy | Capsule 1 | Observed stool | 0.1131 | 9.2% | 0.688 |
| Distal-local low butyrate-associated proxy | Capsule 4 | Arithmetic + reference, target region omitted | 0.1243 | 0.6% | 0.562 |
| Distal-local low butyrate-associated proxy | Capsule 4 | Aitchison + reference, target region omitted | 0.0651 | 47.9% | 0.875 |
| Distal-local low butyrate-associated proxy | Capsule 4 | Observed stool | 0.1242 | 0.7% | 0.562 |

#### S2.7. Empirical channels and state prevalence

Rows of each channel index the regional target state *X* and columns index the thresholded observation *O*.

**Table S5:** Cross-fitted state prevalence and Jeffreys-smoothed empirical channel matrices.

| Task | Measurement/model | $\phi_1$ | $T_{00}$ | $T_{01}$ | $T_{10}$ | $T_{11}$ |
| --- | --- | --- | --- | --- | --- | --- |
| Proximal-local oxygen-tolerant/sulfidogenic proxy | Observed stool | 0.528 | 0.595 | 0.405 | 0.293 | 0.707 |
|  | Population arithmetic stool center | 0.528 | 0.081 | 0.919 | 0.073 | 0.927 |
|  | Population Aitchison stool center | 0.528 | 0.081 | 0.919 | 0.073 | 0.927 |
|  | Arithmetic regions + stool reference | 0.528 | 0.324 | 0.676 | 0.073 | 0.927 |
|  | Aitchison regions + stool reference | 0.528 | 0.595 | 0.405 | 0.415 | 0.585 |
|  | Multi-region CLR ridge | 0.528 | 0.595 | 0.405 | 0.341 | 0.659 |
| Distal-local low butyrate-associated proxy | Observed stool | 0.500 | 0.385 | 0.615 | 0.308 | 0.692 |
|  | Population arithmetic stool center | 0.500 | 0.077 | 0.923 | 0.077 | 0.923 |
|  | Population Aitchison stool center | 0.500 | 0.077 | 0.923 | 0.077 | 0.923 |
|  | Arithmetic regions + stool reference | 0.500 | 0.846 | 0.154 | 0.154 | 0.846 |
|  | Aitchison regions + stool reference | 0.500 | 0.923 | 0.077 | 0.077 | 0.923 |
|  | Multi-region CLR ridge | 0.500 | 0.077 | 0.923 | 0.538 | 0.462 |
| Regional Bacteroidales/Prevotella hotspot proxy | Observed stool | 0.500 | 0.385 | 0.615 | 0.231 | 0.769 |
|  | Population arithmetic stool center | 0.500 | 0.077 | 0.923 | 0.077 | 0.923 |
|  | Population Aitchison stool center | 0.500 | 0.077 | 0.923 | 0.077 | 0.923 |
|  | Arithmetic regions + stool reference | 0.500 | 0.692 | 0.308 | 0.231 | 0.769 |
|  | Aitchison regions + stool reference | 0.500 | 0.410 | 0.590 | 0.308 | 0.692 |
|  | Multi-region CLR ridge | 0.500 | 0.769 | 0.231 | 0.462 | 0.538 |

#### S2.8. Ridge-penalty selection

The prespecified grid was {0.1, 1, 10, 100, 103, 104, 105, 106, 107, 108}. Each path was computed entirely inside the corresponding outer training fold.

**Fig S1:**
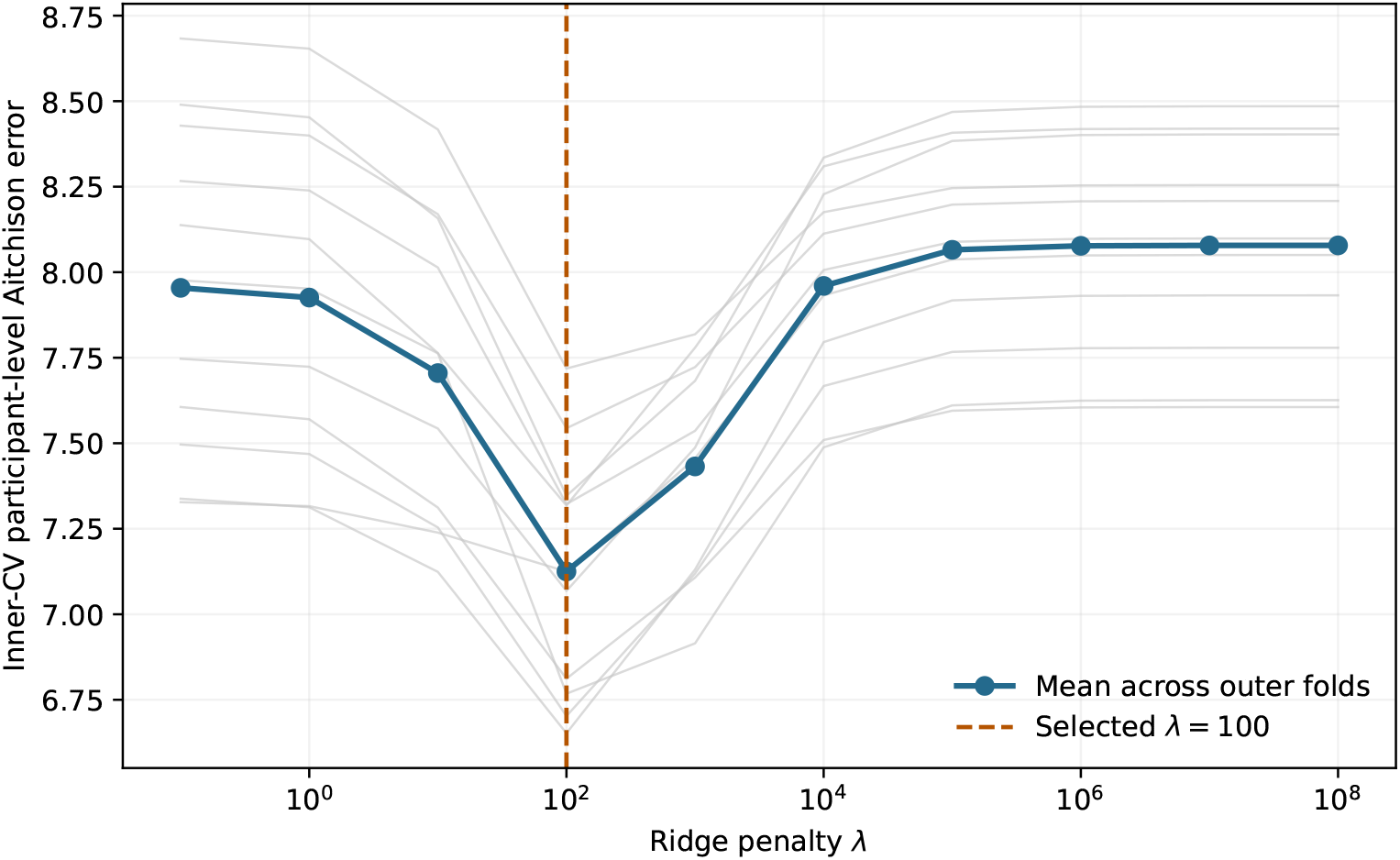
Inner-validation path for multi-region CLR ridge. Gray lines show individual outer folds and the blue line shows their mean. The selected value, *λ* = 100, was the unique minimum in every outer fold and was interior to the grid.

#### S2.9. Complete reconstruction results

| Model | TV [95% CI] | JS [95% CI] | Aitchison [95% CI] |
| --- | --- | --- | --- |
| Multi-region CLR ridge | 0.343 [0.255, 0.440] | 0.367 [0.289, 0.458] | 7.013 [5.593, 8.406] |
| Aitchison regions + stool reference | 0.416 [0.322, 0.516] | 0.430 [0.352, 0.511] | 7.074 [6.297, 7.921] |
| Equal arithmetic regional mean | 0.538 [0.474, 0.626] | 0.565 [0.511, 0.632] | 7.578 [6.869, 8.296] |
| Learned arithmetic regional mixture | 0.529 [0.466, 0.612] | 0.557 [0.501, 0.628] | 7.647 [6.908, 8.528] |
| Equal Aitchison regional center | 0.577 [0.510, 0.659] | 0.599 [0.536, 0.668] | 7.752 [6.950, 8.693] |
| Capsule 4 | 0.510 [0.421, 0.614] | 0.541 [0.444, 0.631] | 8.004 [6.293, 9.881] |
| Learned Aitchison regional mixture | 0.565 [0.496, 0.650] | 0.592 [0.525, 0.667] | 8.019 [6.942, 9.237] |
| Population Aitchison stool center | 0.332 [0.257, 0.417] | 0.369 [0.298, 0.454] | 8.039 [6.582, 9.416] |
| Capsule 3/4 arithmetic mean | 0.552 [0.477, 0.641] | 0.587 [0.527, 0.655] | 8.351 [7.270, 9.776] |
| Capsule 2 | 0.594 [0.500, 0.695] | 0.620 [0.538, 0.711] | 8.644 [7.300, 10.040] |
| Capsule 1 | 0.576 [0.494, 0.662] | 0.588 [0.510, 0.675] | 9.371 [7.696, 11.136] |
| Capsule 3 | 0.633 [0.561, 0.714] | 0.669 [0.618, 0.723] | 9.472 [8.130, 10.832] |
| Arithmetic regions + stool reference | 0.319 [0.264, 0.388] | 0.374 [0.326, 0.431] | 9.721 [8.368, 11.092] |
| Population arithmetic stool center | 0.316 [0.256, 0.385] | 0.374 [0.322, 0.434] | 10.061 [8.644, 11.512] |

#### S2.10. Prespecified paired tests

Positive improvement means model A had lower participant-level error than model B. Holm adjustment was performed separately within total-variation and Aitchison families.

| Metric | Model A | Model B | Mean improvement | Exact *P* | Holm *P* |
| --- | --- | --- | --- | --- | --- |
| TV | Arithmetic regions + stool reference | Population arithmetic stool center | -0.003 | 0.6426 | 0.7520 |
| TV | Aitchison regions + stool reference | Population arithmetic stool center | -0.099 | 0.0088 | 0.0264 |
| TV | Multi-region CLR ridge | Population arithmetic stool center | -0.027 | 0.3760 | 0.7520 |
| TV | Arithmetic regions + stool reference | Equal arithmetic regional mean | 0.219 | 0.0020 | 0.0078 |
| TV | Aitchison regions + stool reference | Equal Aitchison regional center | 0.161 | 0.0010 | 0.0049 |
| Aitchison | Arithmetic regions + stool reference | Population Aitchison stool center | -1.681 | 0.0010 | 0.0049 |
| Aitchison | Aitchison regions + stool reference | Population Aitchison stool center | 0.965 | 0.0947 | 0.2432 |
| Aitchison | Multi-region CLR ridge | Population Aitchison stool center | 1.026 | 0.0811 | 0.2432 |
| Aitchison | Arithmetic regions + stool reference | Equal arithmetic regional mean | -2.143 | 0.0264 | 0.1055 |
| Aitchison | Aitchison regions + stool reference | Equal Aitchison regional center | 0.678 | 0.1953 | 0.2432 |

#### S2.11. Crossed guild-resolution and zero-replacement sensitivity

Positive paired differences and positive relative reductions favor the regional-plus-reference model. Intervals are participant-bootstrap intervals; exact *P* values are unadjusted descriptive sensitivity tests.

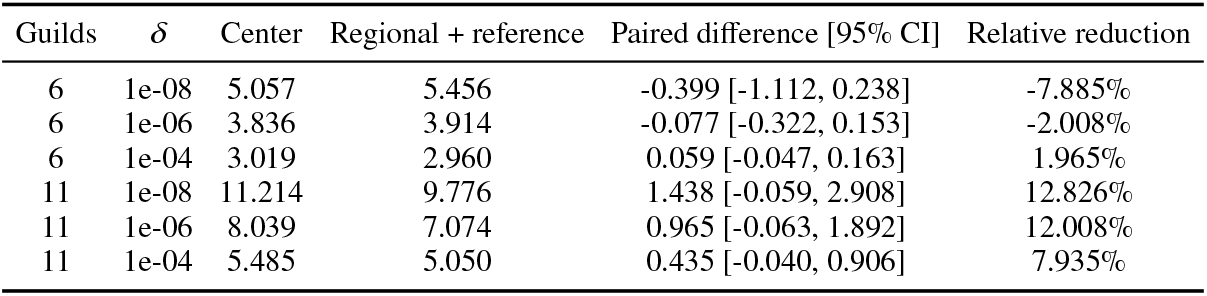

| Guilds | $\delta$ | Center | Regional + reference | Paired difference [95% CI] | Relative reduction |
| --- | --- | --- | --- | --- | --- |
| 6 | 1e-08 | 5.057 | 5.456 | -0.399 [-1.112, 0.238] | -7.885% |
| 6 | 1e-06 | 3.836 | 3.914 | -0.077 [-0.322, 0.153] | -2.008% |
| 6 | 1e-04 | 3.019 | 2.960 | 0.059 [-0.047, 0.163] | 1.965% |
| 11 | 1e-08 | 11.214 | 9.776 | 1.438 [-0.059, 2.908] | 12.826% |
| 11 | 1e-06 | 8.039 | 7.074 | 0.965 [-0.063, 1.892] | 12.008% |
| 11 | 1e-04 | 5.485 | 5.050 | 0.435 [-0.040, 0.906] | 7.935% |

#### S2.12. Decision-layer sensitivity at six-guild resolution

The proxy-state and observed-stool scores are exact amalgamation equivalents across K=11 and K=6. Models were refitted at K=6; values below are point-estimate mean MIDL-curve calibration errors across the three tasks.

| Model | $K=11$ | $K=6$ |
| --- | --- | --- |
| Population Aitchison stool center | 0.0053 | 0.0053 |
| Arithmetic regions + stool reference | 0.0298 | 0.0318 |
| Aitchison regions + stool reference | 0.0327 | 0.0144 |
| Multi-region CLR ridge | 0.0143 | 0.0033 |

#### S2.13. Additional figures

**Fig S2:**
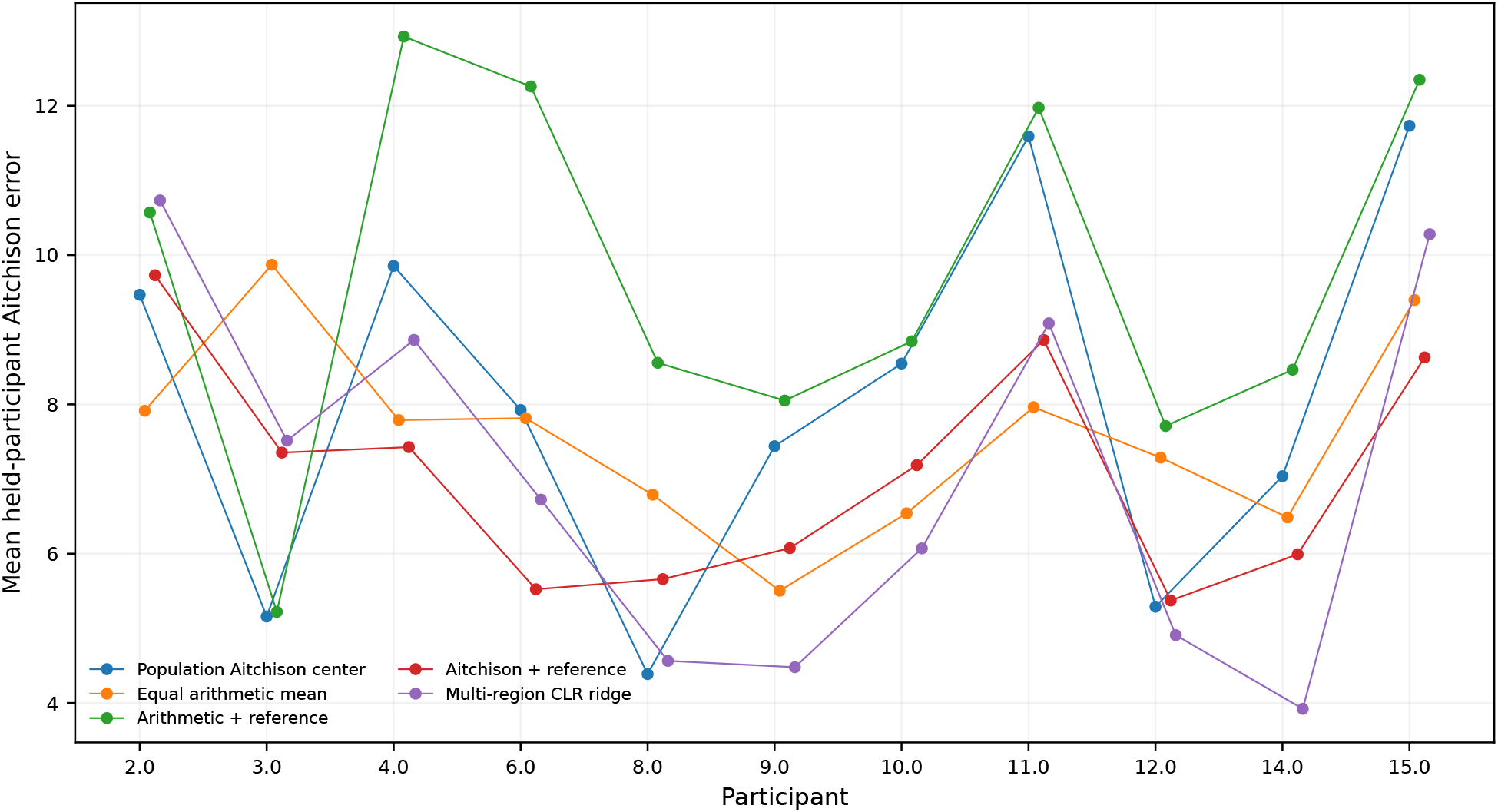
Held-participant Aitchison errors for selected models. Lines connect model-specific observations only for visualization; inference used paired participant-level differences.

**Fig S3:**
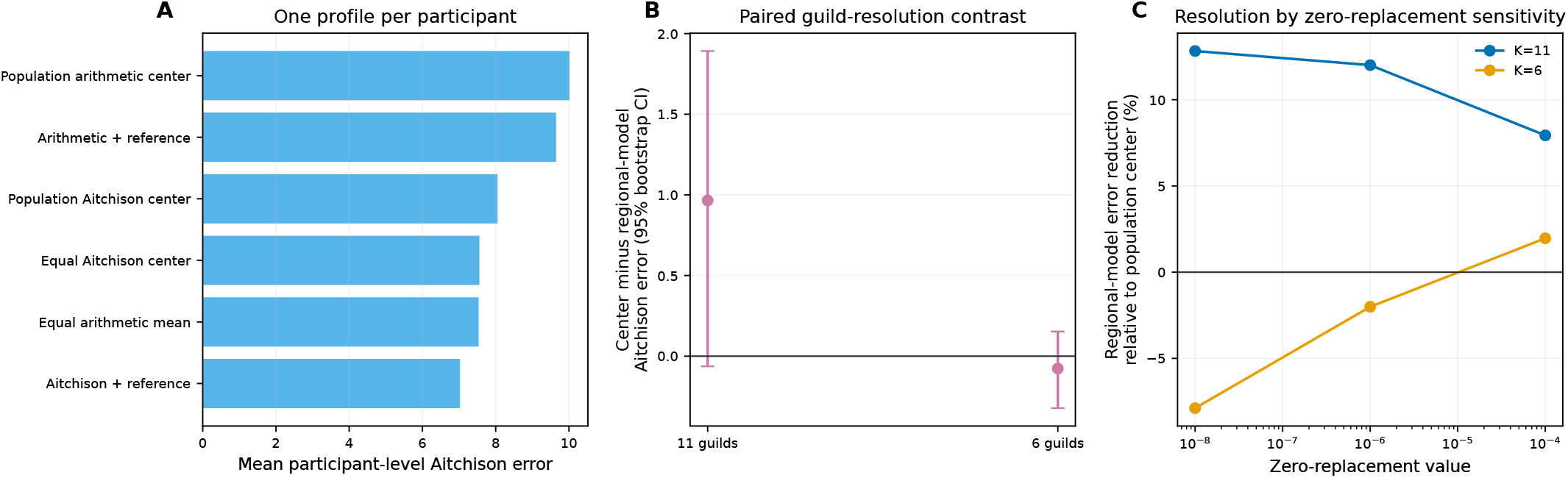
Representation and numerical sensitivities. (A) Reconstruction after averaging repeated events to one profile per participant. (B) Paired participant-level contrast between the population Aitchison center and the regional-plus-reference model at 11- and 6-guild resolution; positive values favor the regional model. (C) Relative error reduction of the regional model across crossed guild-resolution and zero-replacement choices.

## Notes

### Competing Interest Statement

The authors have declared no competing interest.

